# Epigenomic profiling of neuroblastoma cell lines

**DOI:** 10.1101/829754

**Authors:** Kristen Upton, Apexa Modi, Khushbu Patel, Karina L. Conkrite, Robyn T. Sussman, Gregory P. Way, Rebecca N. Adams, Gregory I. Sacks, Paolo Fortina, Sharon J. Diskin, John M. Maris, Jo Lynne Rokita

## Abstract

Understanding the aberrant transcriptional landscape of neuroblastoma is necessary to provide insight to the underlying influences of the initiation, progression and persistence of this developmental cancer. Here, we present chromatin immunoprecipitation sequencing (ChIP-Seq) data for the oncogenic transcription factors, MYCN and MYC, as well as regulatory histone marks H3K4me1, H3K4me3, H3K27Ac, and H3K27me3 in ten commonly used human neuroblastoma-derived cell line models. In addition, for all of the profiled cell lines we provide ATAC-Seq as a measure of open chromatin. We validate specificity of global MYCN occupancy in MYCN amplified cell lines and functional redundancy of MYC occupancy in MYCN non-amplified cell lines. Finally, we show with H3K27Ac ChIP-Seq that these cell lines retain expression of key neuroblastoma super-enhancers (SE). We anticipate this dataset, coupled with available transcriptomic profiling on the same cell lines, will enable the discovery of novel gene regulatory mechanisms in neuroblastoma.

## Background & Summary

An estimated 15,780 children in the United States will be diagnosed with cancer in 2019 (ACCO 2016). While 80% of pediatric cancer patients overcome this disease, 20% of children do not survive, and survivors often have multiple side effects of therapy (ACCO 2016). Neuroblastoma accounts for more than 7% of malignancies in patients under 15 years of age and approximately 12% of all pediatric cancer-related deaths (for review (Matthay et al. 2016)). Neuroblastoma shows wide phenotypic variability, with tumors arising in children diagnosed under the age of 18 months often spontaneously regressing with little or no treatment, but patients diagnosed at an older age or with unfavorable genomic features often showing a relentlessly progressive and widely metastatic disease pattern despite intensive, multimodal therapy (for review see (Matthay et al. 2016; Maris 2010)). Ninety-eight percent of low-risk neuroblastoma disease are currently cured (Twist et al. 2019), however, the survival rate for patients with high-risk neuroblastoma remains less than 50% (Pinto et al. 2015). Relapsed high-risk neuroblastoma is typically incurable (Simon et al. 2011; Ambros et al. 2009), and thus these children require improved therapeutic options.

A major prognostic factor predicting the severity, risk, and inferior outcome for neuroblastoma patients is amplification of the proto-oncogene *MYCN. MYCN* amplification occurs in nearly 20% of all neuroblastomas, and approximately 50% of patients with high-risk disease (Gherardi et al. 2013; Rickman, Schulte, and Eilers 2018). It is a truncal genomic event, and typical stable across the spectrum of therapy and disease recurrence. MYCN, along with structural and binding homologues MYC and MYCL, are members of the MYC transcription factor family (Pistoia et al. 2012) and have been implicated in transcriptional regulation of proteins involved in cell growth (Coller et al. 2000), proliferation (Ji et al. 2011), and ribosome biogenesis (Ji et al. 2011). Mounting evidence has also indicated that MYCN and MYC are functionally redundant (He et al. 2013; Malynn et al. 2000; Chappell and Dalton 2013). However, the protein expression of MYC and MYCN appears to be mutually exclusive. For example, neuroblastoma tumors with *MYCN* amplification typically lack or have low *MYC* mRNA expression (Rickman, Schulte, and Eilers 2018). The strong influence of MYCN on the progression and metastasis of neuroblastoma makes it a key target for therapy, but due to its global transcriptional activity, it is necessary to develop a better understanding of which of its gene targets directly influence oncogenesis.

To better understand the regulatory effects of MYC family proteins in neuroblastoma, we performed ChIP-Seq data for MYCN in six neuroblastoma cell lines with *MYCN* amplification, MYC in four neuroblastoma cell lines without *MYCN* amplification, and H3K27Ac, H3K27me3, H3K4me1, and H3K4me3 histone modifications along with ATAC-Seq in all ten neuroblastoma cell lines (with ATAC data in four additional lines also reported here). All of the cell lines here also have RNA sequencing data freely available (Harenza et al. 2017).

## Methods

Online Table 1 summarizes which assays were performed for each cell line, and an overview of the workflow is shown in Figure 1.

**Figure 1.**
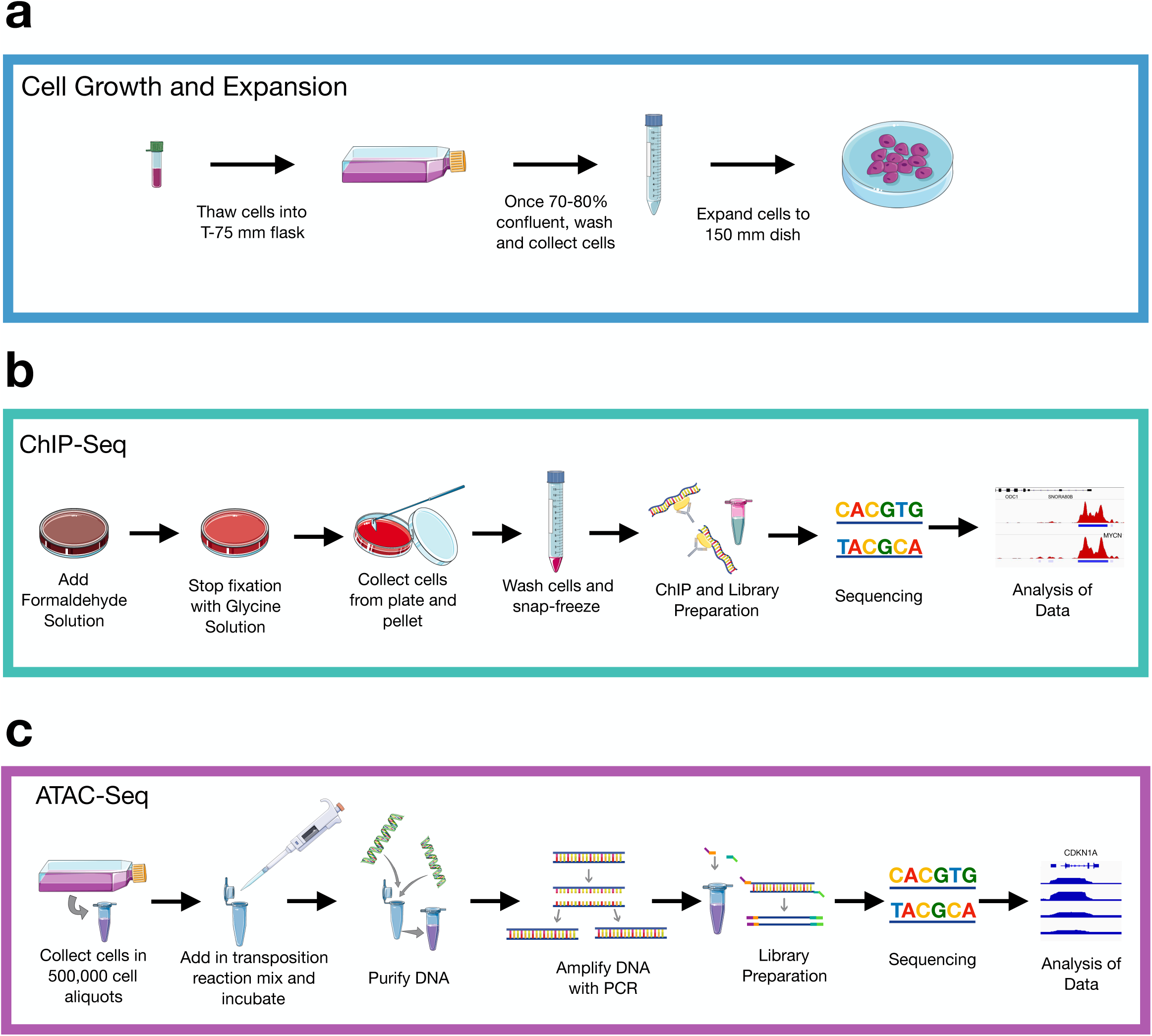
Experimental workflow. (**A**) Cells were thawed, grown, and expanded until 70-80% confluency in a 150 mm dish. (**B**) For ChIP-Seq, cells were fixed, collected, and frozen (N = 1 biological replicate per cell line). Libraries were prepared, sequenced, and data analyzed. (**C**) For ATAC-Seq (n = 14 samples with n = 2 biological replicates), cells were incubated in a transposition reaction, DNA was purified, and amplified with limited PCR. Libraries were prepared, sequenced, and analyzed. Diagram was created using Servier Medical ART (https://smart.servier.com/).

### Cell Growth and Expansion

The cell lines used to collect this data were obtained from multiple sources: the Children’s Oncology Group (COG) Cell Culture and Xenograft Repository at Texas Tech University Health Sciences Center (www.cccells.org), the American Type Culture Collection (Manassas, MA), or the Children’s Hospital of Philadelphia (CHOP) cell line bank. All of the cell growth and preparations were done at CHOP. The neuroblastoma cell lines were cultured using media (Metadata Table 1) and methods as previously described (Harenza et al. 2017). Briefly, cells were thawed by floating in a 37°C water bath for 2-3 minutes. Cells were then added to a 15 mL conical tube, containing 5 mL of the appropriate growth media, and centrifuged at 300 *xg* for 5 minutes at room temperature (RT). Media was then carefully aspirated off, and the pelleted cells were resuspended in 1 mL of media before being transferred to a 75 mm^2^ flask containing 10 mL of growth media. Cells were incubated at 37°C with a 5.0% CO_2_ concentration. When cells reach 70-80% confluency, media was aspirated off and cells were gently washed with 1X PBS. Following aspiration of the PBS, 3 mL of the appropriate detachment solution (noted in Metadata Table 1) was added and the flask was incubated at 37°C for 2-5 minutes. Cells were then gathered by tilting the plate at a 45° angle and washing with at least 4 mL of the appropriate growth media, and transferred to a 15 mL conical. After centrifugation for 5 minutes at 300 *x g*. Media was aspirated off, and the pellet was resuspended in 1 mL of growth media and transferred to a 150 mm cell culture dish containing 19 mL of growth media. Cells were incubated at 37°C with a 5.0% CO_2_ concentration until reaching 70-80% confluency. Necessary materials and reagents are listed in Online Table 2.

### ChIP-Seq Protocol

The ChIP-Seq Protocol is separated into four sections: Cell Fixation, Chromatin Immunoprecipitation (ChIP) and Library Preparation, Library Sequencing, and ChIP-Seq Analysis. Of note, the MYCN ChIP-Seq for Kelly and NGP cell lines was performed using a varied procedure and is noted in a separate section within this protocol. Necessary materials and reagents are listed in Online Table 2.

#### Cell Fixation

Cells were grown as described in Cell Growth and Expansion section of protocol to 70-80% confluence in 150 mm tissue culture plates in 20 mL of media. The Formaldehyde solution (Online Table 2) was freshly prepared. Cells were removed from incubation and 1/10th of the growth media volume of the Formaldehyde Solution was added to the existing media in the plate (i.e. if the current volume of the plate is 20 mL of media, 2 mL of Formaldehyde Solution would be added). The solution was gently swirled, and then rocked at RT for 15 minutes. To stop the fixation, 1/20th the current volume of the Glycine Solution (Online Table 2) was added to the plate (i.e. if the current volume in the plate is 22 mL then 1.1 mL of Glycine Solution should be added). The plate was gently swirled to mix, and then allowed to sit at RT for 5 minutes. Following this incubation, a cell scraper was used to collect the cells, and then all cells and solution were transferred to a 50 mL conical on ice. From this point forward, all samples were kept on ice. The 50 mL conical was centrifuged at 800 *x g* at 4°C for 10 minutes to pellet the cells. Supernatant was removed and discarded, and the cells were resuspended with 10 mL of chilled, sterile PBS. Centrifugation of the tube at 800 *x g* at 4°C for 10 minutes was repeated. The supernatant was removed and discarded, and the cells were resuspended with 10 mL of chilled, sterile PBS with 100 uL of PMSF. The tube was centrifuged at 800 *x g* at 4°C for 10 minutes, the supernatant was removed, and then the cells were snap frozen on dry ice and stored at -80°C. The cells were then shipped to Active Motif on dry ice following the instructions listed at on the Sample Submission Form, downloaded from www.activemotif.com/sample-submission.

#### ChIP and Library Preparation by Active Motif

Chromatin immunoprecipitation was completed by Active Motif. Full methods are proprietary. Chromatin was isolated using a lysis buffer and membranes were disrupted with a dounce homogenizer. The lysates were then sonicated with Active Motif’s EpiShear probe sonicator (#53051) and cooled sonication platform (#53080) to an average fragment length 300-500 bp. A portion of the sample was collected as the Input DNA, treated with RNase, proteinase K, and incubated to reverse crosslinking. The DNA was then collected by ethanol precipitation. The Input DNA was resuspended and concentration was quantified by a NanoDrop spectrophotometer. Extrapolation of this concentration to the original chromatin volume allowed for quantitation of the total chromatin yield. Aliquots of the fixed chromatin were used in the immunoprecipitation were precleared with protein A agarose beads (Invitrogen, #15918014). Genomic DNA regions of interest were isolated using specific ChIP antibodies (Online Table 2). Antibody DNA complexes were isolated using additional protein A agarose beads, and the crosslinked DNA, antibody, and bead complexes were washed. The cross-linked DNA was eluted from the beads with SDS buffer, and subjected to RNase and proteinase K treatment. Reverse crosslinking was done in an overnight incubation at 65°C, and ChIP DNA was purified with a phenol-chloroform extraction and ethanol precipitation.

Illumina sequencing libraries were prepared from the ChIP and Input DNAs using the standard consecutive enzymatic steps of end-polishing, dA-addition, and adaptor ligation using Active Motif’s custom liquid handling robotics pipeline. Samplers were amplified with a 15 cycle PCR amplification and then quantified before being shipped to the Jefferson Cancer Genomics Laboratory at the Kimmel Cancer Center for sequencing.

#### MYCN ChIP-Seq: Kelly and NGP Cell Lines

Chromatin immunoprecipitation was performed on adherent cells as described in Bosse et al., 2017(Bosse et al. 2017). Of note, a different MYCN antibody was used than listed in Bosse et al., 2017 (Santa Cruz B8.4B, sc-53993). Cells were grown as described in Cell Growth and Expansion section of protocol to 70-80% confluence in 150 mm tissue culture plates in 20 mL of media. To the existing media, 415 mL of 37% formaldehyde (final concentration of 0.75%) was added, and rocked for 10 minutes at RT. To this, 1.5 mL of 2.5 M glycine (Online Table 2) (final concentration of 0.18 M) was added to inactivate the formaldehyde, and the plate was rocked for an additional 5 min. Cells were lysed with a volume of FA Lysis Buffer (Online Table 2) equivalent to 5 pellet volumes. Beads were washed 3 times in ChIP Wash Buffer (Online Table 2) and one time with Final Wash Buffer (Online Table 2). Libraries were constructed using NEB Ultra Kit following manufacturer’s instructions. Libraries were sequenced as single-end, 50 bp reads on a MiSeq to a depth of ∼50 M reads by the Children’s Hospital of Philadelphia Nucleic Acid and PCR Core.

#### ChIP Library Sequencing for ChIP

Sequencing was conducted by the Jefferson Cancer Genomics Laboratory at the Kimmel Cancer Center. Samples were quality control tested using an Agilent High Sensitivity Screen Tape to determine average fragment length. The concentration of each library was measured using a High Sensitivity Qubit Quantification kit, and samples were diluted to an appropriate amount for the loading protocol (4 nM or less). Samples were normalized to the same nanomolar concentration, and libraries were pooled together in equal amounts. Samples were diluted to 1.51 pM in Low EDTA TE Buffer. Samples were then sequenced as single-end, 70 bp reads to an average depth of ∼30 M reads on a NextSeq 500.

### ATAC-Seq Protocol

The following ATAC-Seq protocol was adapted from Buenrostro, et. al, 2015(Buenrostro et al. 2015). This protocol consists of four parts: Cell Preparation, Transposition Reaction and Purification, PCR Amplification, qPCR, and Library Preparation. Primer 1 and Primer 2 were custom synthesized by Integrated DNA Technologies (IDT), using sequences provided in Buenrostro, et. al, 2015. Note: ATAC-Seq for NB-69 and NGP was performed using a slightly varied procedure and is noted in a separate section.

#### Cell Preparation

Cells were grown as described in Cell Growth and Expansion section of protocol to 70-80% confluence in a 75 mm^2^ tissue culture flasks in 10 mL of media. Following detachment and pelleting, cells were resuspended in 1.0 mL of the appropriate growth media. Cells were triturated until they were in a homogenous single-cell suspension. Using an automated cell counter, the volume for 500,000 cells was determined and aliquoted into a sterile 1.5 mL Eppendorf tube containing 500 μL of sterile 1X PBS. Cells were centrifuged at 500 *x g* for 5 minutes at 4°C. The supernatant was carefully aspirated, and the cells were resuspended in 500 mL of sterile 1X PBS. Centrifugation was repeated and cells were resuspended in 500 mL of cold lysis buffer by gently pipetting up and down, and then immediately centrifuged at 500 *x g* for 10 minutes at 4°C. The supernatant was carefully removed and discarded. The pellet was immediately resuspend in 50 μL of nuclease free water by gently pipetting up and down, and the protocol immediately continued on to Transposition Reaction and Purification section.

#### Transposition Reaction and Purification

The pellet was placed on ice. The following reagents were prepared and combined: transposition reaction mix (25 μL TD (2X reaction buffer from Nextera Kit), 2.5 μL TDE1 (Nextera Tn5 Transposase from Nextera Kit), 17.5 μL nuclease-free water, and 5.0 μL of resuspended DNA/protein from the final step in *Cell Preparation* (resuspended pellet in 50 μL of nuclease free water). The transposition reaction was incubated in a thermocycler at 37°C for 30-35 minutes. The reaction was immediately purified using Qiagen MinElute PCR Purification Kit, and the transposed DNA was eluted in 10.5 μL of elution buffer (Buffer EB from the MinElute Kit consisting of 10 mM Tris-Cl (pH 8)). The eppendorf tube containing purified DNA was parafilmed, and stored at -20°C. *NOTE:* This can act as a good stopping point, however these DNA fragments are not PCR amplifiable if melted at this point.

#### PCR Amplification

Primer sequences are shown in Online Table 3. To amplify the Transposed DNA, the following were combined into a 0.2 mL PCR tube: 10 μL transposed DNA, 10 μL nuclease-free H_2_O, 2.5 μL 25 mM PCR Primer 1 (Ad1), 2.5 μL 25 mM Barcoded PCR Primer 2 (Ad2.X, X being the unique number of samples), and 25 μL NEBNext High-Fidelity 2X PCR Master Mix. The thermal cycle was as follows:

|  |  |
| --- | --- |
| 1 cycle: | 5 min 72°C |
|  | 30 sec 98°C |
| 5 cycles: | 10 sec 98°C |
|  | 30 sec 63°C |
|  | 1 min 72°C |

The five minute extension in the first cycle is critical to allow extension on both ends of the primer after transposition, thereby generating amplifiable fragments. This ensures that downstream quantitative PCR (qPCR) quantitation will not change the complexity of the original library.

#### qPCR

To reduce the GC and size bias in PCR, the appropriate number of PCR cycles (N) was determined using qPCR, allowing us to stop prior to saturation. The samples were kept in the thermocycler following the PCR Amplification reaction, and the qPCR side reaction was run. In a 0.2 mL PCR tube the following were added: 5 μL of DNA PCR amplified DNA, 2 μL of nuclease free H--_2_O, 1 μL of 6.25 mM Custom Nextera PCR Primer (Ad1), 1 μL of 6.25 mM Custom Nextera PCR Primer 2 (Ad2.X), 1 μL 9X SYBR Green I, and 5 μL NEBNext High-Fidelity 2X PCR Master Mix. This sample was run in the qPCR instrument with the following cycles:

|  |  |
| --- | --- |
| 1 cycle: | 30 sec 98°C |
| 20 cycles: | 10 sec 98°C |
|  | 30 sec 63°C |
|  | 1 min 72°C |

To calculate the additional number of cycles needed, a linear plot of Rn versus cycle was generated. This determined the cycle number (N) that corresponds to one-third of the maximum fluorescent intensity.

The remaining 45 mL PCR reaction was run to the cycle number (N) determined by qPCR. Cycles are as follows:

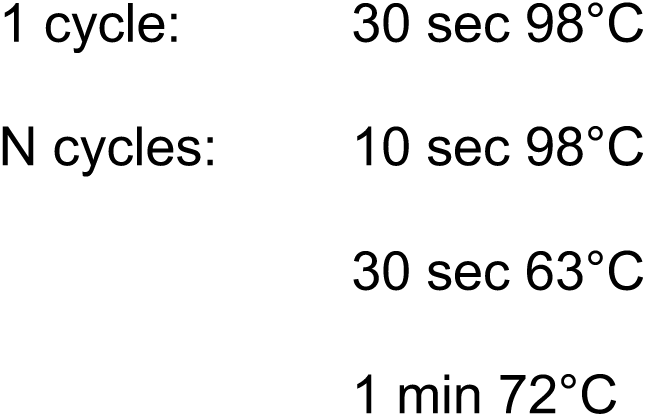

|  |  |
| --- | --- |
| 1 cycle: | 30 sec 98°C |
| N cycles: | 10 sec 98°C |
|  | 30 sec 63°C |
|  | 1 min 72°C |

The amplified library was purified using Qiagen MinElute PCR Purification Kit after the additional PCR. The purified library was eluted in 20 μL of elution buffer (Buffer EB from the MinElute Kit consisting of 10 mM Tris-Cl (pH 8)). It is important to make sure that the column is dry prior to adding elution buffer to avoid ethanol contamination of final library. The amplified library was purified using AMPure XP beads at a 1.8x ratio to get rid of adapter dimers, using 80% ethanol for the wash steps. Sample was eluted in 50 μL of nuclease free H_2_O. The concentration of the DNA eluted from the column should be about 30 nM.

#### Library Preparation

The quality of the purified libraries was assessed using a Bioanalyzer High-Sensitivity DNA Analysis kit (Agilent). If libraries contained predominant peaks around 1000 bp, SPRI beads were used to remove these fragments. This was accomplished by first, with a new vial of SPRI beads, performing size selection with various ratios to ensure larger peaks are removed. For example, ratios could include 0.4X, 0.45X, 0.5X. Choose the ratio that removes 1000 bp fragments, but leaves 800 bp fragments. Libraries were eluted in 20 μL of nuclease-free water, and sequenced as described below.

#### Sequencing for ATAC-Seq by Beijing Genomics International (BGI)

Sequencing was conducted by Beijing Genomics International at the Children’s Hospital of Philadelphia. Samples were quality control tested using an Agilent High Sensitivity Screen Tape to confirm average fragment sizes were ∼180, 380, 580, 780, and 980 bp. The concentration of each library was measured using a High Sensitivity Qubit Quantification kit, to ensure they were 5.5 nM. Samples were normalized and libraries were pooled together in equal amounts. Samples were then sequenced as paired-ends, 100 bp to an average depth of 80M reads on a HiSeq 2500.

#### ATAC-Seq NB-69 and NGP cell lines via Active Motif

Cells were grown as described in Cell Growth and Expansion section of protocol to 70-80% confluence in a 75 mm^2^ tissue culture flasks in 10 mL of media. Following detachment and pelleting, cells were resuspended in 1.0 mL of the appropriate growth media. Cells were triturated into a homogenous single-cell suspension. Using an automated cell counter, the volume for 100,000 cells was determined, and aliquoted into a sterile 1.5 mL eppendorf tube containing 500 μL of sterile 1X PBS. Cells were then centrifuged at 500 *x g* for 5 minutes at 4°C. The supernatant was carefully aspirated off, and the cells were resuspended in 500 μL of growth media with 5% DMSO. The sample was transferred to a 1.7 mL microfuge tube on ice. Cells were frozen with a slow cooling to minimize cell lysis. Samples were shipped on dry ice to Active Motif (1914 Palomar Oaks Way, Ste 150, Carlsbad, CA 92008) following the instructions listed at on the Sample Submission Form, downloaded from www.activemotif.com/sample-submission. Samples were prepared and sequenced following Active Motif’s ATAC-Seq proprietary protocol. Cells were thawed in a 37°C water bath, pelleted, washed with cold PBS, and tagmented as previously described (Buenrostro et al. 2015), with some modifications based on (Corces et al. 2017). Cell pellets were resuspended in lysis buffer, pelleted, and tagmented using the enzyme buffer provided in the Nextera Library Prep Kit (Illumina). Tagmented DNA was then purified using the MinElute PCR purification kit (Qiagen), amplified with 10 cycles of PCR, and purified using Agencourt AMPure SPRI beads (Beckman Coulter). The resulting material was quantified using the KAPA Library Quantification Kit for Illumina platforms (KAPA Biosystems) and sequenced with PE42 sequencing on the NextSeq 500 sequencer (Illumina).

### ChIP-Seq Data Analysis

FASTQ quality was assessed using FastQC v0.11.4 (http://www.bioinformatics.babraham.ac.uk/projects/fastqc/) and sequences were adapter- and quality-trimmed using Trim Galore v.0.4.0 (https://github.com/FelixKrueger/TrimGalore) and CutAdapt v.1.12 (Krueger n.d.; Lindgreen 2012). Since multiple sequencers were used, FASTQ phred sequencing scores (Ewing et al. 1998) were calculated using a perl script (https://raw.githubusercontent.com/douglasgscofield/bioinfo/master/scripts/phredDetector.pl).

This value was used as input into the alignment algorithm. The bwa v.0.7.12 samse (Li and Durbin 2009) was used to align the reads to hg19 reference genome and Picard tools v.2.17.9-SNAPSHOT (Alec Wysoker KT, Mike McCowan, Nils Homer, Tim Fennell, n.d.) was used to remove duplicates. Fragment sizes were estimated using MaSC 1.2.1 (Ramachandran et al. 2013) and these values were used as input into MACS2 v.2.1.1 (Zhang et al. 2008) for narrow peak calling (transcription factors) or broad peak calling (histone marks). Broad peaks were called significant using a q-value (minimum False Discovery Rate) cut off of 0.05 and narrow peaks at a q-value cutoff of 0.1. Results were returned in units of signal per million reads to get normalized peak values. Finally, repetitive centromeric, telomeric and satellite regions known to have low sequencing confidence were removed using the blacklisted regions defined by UCSC.

### ATAC-Seq Data Analysis

Samples were quality-controlled and trimmed as described in Chip-Seq Analysis. FASTQ files were aligned using bwa aln for BGI samples (100 bp reads) and bwa mem for Active Motif samples (42 bp reads). Reads with mapping quality < 10 were discarded. Biological duplicate BAMs were merged using Picard v.2.17.9-SNAPSHOT. Broad peaks were called using --extsize 200, --shift 100, and using a q-value (minimum False Discovery Rate) cut off of 0.1. Results were returned in units of signal per million reads to get normalized peak values. Finally, repetitive centromeric, telomeric and satellite regions known to have low sequencing confidence were removed using the blacklisted regions defined by UCSC.

### ChIP-Seq Quality Control Metrics

We investigated three metrics to assess ChIP-seq quality. To calculate enrichment of reads within peaks we determined the FRiP score using deeptools2 (Ramírez et al. 2016). The FRiP score is defined as the fraction of reads that fall within a peak divided by the total number of reads. To measure read enrichment independent of peak calling we calculated the NSC (normalized strand cross-correlation) and the RSC (relative strand cross-correlation) using phantompeakqualtools (Landt et al. 2012; Kharchenko, Tolstorukov, and Park 2008)as part of the ENCODE ChIP-seq processing pipeline. All ChIP-Seq data passed quality control and results are reported in Online Table 4.

### Super-enhancer Calling and Comparison

Super-enhancers (SEs) were called from H3K27Ac BAM files using the default parameters of LILY (https://github.com/BoevaLab/LILY), which includes correction for copy number variation inherently present in cancer samples. Enhancers were classified into SEs, enhancers, and promoters and annotated using Homer v4.10.4. Scripts to run LILY can be found on Github (https://github.com/marislab/epigenomics-data-descriptor). SEs were also called from H3K27Ac MACS2 peaks using ROSE v.0.1 (https://bitbucket.org/young_computation/rose/src/master/) using default parameters and annotated using Homer v4.10.4. SEs which overlapped with the MYCN locus (hg19, chr2:16080683-16087129) were removed from the analysis. SE genes which we annotated as transcription factors (Lambert et al. 2018) were used for comparison to two literature studies (Boeva et al. 2017; van Groningen et al. 2017).

### Heatmap Preparation

The 5,000 most significant (sorted by highest -log_10_(p-value) and -log_10_(q-value)) MYCN peaks for each of the five MYCN amplified cell line were intersected using bedtools. Heatmaps were generated for regions +/-4 kb from the transcription start site (TSS) for the 5,046 peaks common to at least four MYCN amplified cell lines. Heatmaps were created for LA-N-5 and NB-69 at loci annotated as enhancers, SEs, and promoters-TSS by LILY. All ChIP-seq heatmaps were created using deepTools 3.2.0 package plotHeatmap tool (Ramírez et al. 2016). The code and parameters used to generate heatmaps can be found on GitHub (https://github.com/marislab/epigenomics-data-descriptor).

### Cell Line Authentication

All cell lines were STR-authenticated by Guardian Forensic Sciences (Abington, PA) using the GenePrint 24 (Promega, #B1870).

## Data Records

Raw, concatenated FASTQ files and processed BIGWIG files for all sequencing data were deposited into the Gene Expression Omnibus (GEO) under SuperSeries Accession Number GSE138315. MYCN and MYC ChIP-Seq data for the Kelly and NGP cell lines were deposited into GEO under Accession Number GSE94782, all other MYCN and MYC ChIP-Seq were deposited under Accession Number GSE138295, histone ChIP-Seq data were deposited under Accession Number GSE138314, and ATAC-Seq data were deposited under Accession Number GSE138293.

## Technical Validation

Prior to selecting cell lines for MYCN and MYC profiling, we assessed RNA expression (Figure 2A-B) and protein expression (Figure 2C-D) across a subset of neuroblastoma cell lines. NB-LS, while *MYCN* non-amplified, has substantial MYCN RNA and protein expression (Cohn et al. 1990), but was not chosen, as we restricted MYCN ChIP-Seq to *MYCN* amplified cell lines plus one negative control. SK-N-BE(2)-C, a *MYCN* amplified cell line, showed high *MYCN* mRNA expression, but surprisingly low protein expression, and thus was excluded. The remaining cell lines had concordant *MYCN* and *MYC* mRNA and protein expression, thus, COG-N-415, KELLY, NB-1643, LA-N-5, and NGP were chosen for MYCN ChIP-Seq while NB-69, SK-N-AS, and SK-N-SH were chosen for MYC ChIP-Seq. As additional controls, we performed MYCN ChIP-Seq in the *MYCN* non-amplified line NB-69, and MYC ChIP-Seq on the *MYCN* amplified cell line KELLY. To validate the MYCN and MYC ChIP-Seq antibodies, we first intersected loci bound by MYCN in two or more cell lines and of the 157 MYCN transcriptional targets previously reported using ChIP-on-ChIP (Valentijn et al. 2012), found 139 loci occupied by the MYCN via ChIP-Seq (Figure 2E). Next, we integrated the top 5,000 MYCN peaks from each *MYCN* amplified cell line. We generated heatmaps for the peaks (1,330) which overlapped in all five cell lines (as defined in ***Heatmap Preparation***) and depict occupancy of MYCN (Figure 2F) and MYC (Figure 2G) at these sites. As expected, the MYCN amplified cell lines COG-N-415, KELLY, NB-1643, LA-N-5, and NGP show similar binding profiles, while the negative control MYCN non-amplified line NB-69 depicted an absence of binding for MYCN at the same loci. We observed MYC bound to the same loci in the MYCN non-amplified cell lines, SK-N-AS, SK-N-SH, and NB-69 as well as the *MYCN* amplified and low MYC-expressing line KELLY (Figure 2G), supporting the notion of redundant functionality of MYC family protein members.

**Figure 2.**
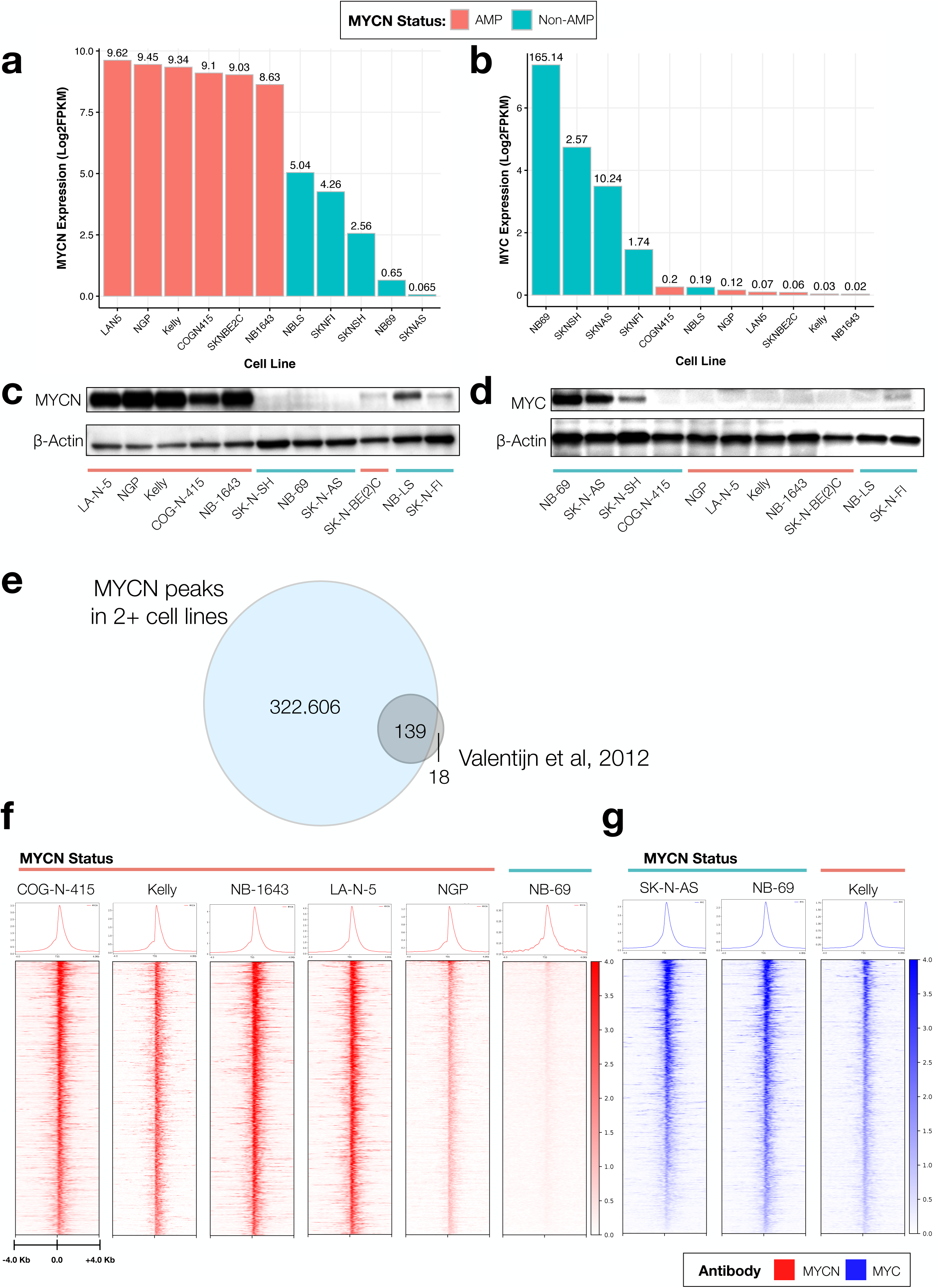
Comparison of MYCN and MYC binding based on MYCN amplification status. The log_2_ FPKM mRNA expression of *MYCN* (**A**) and *MYC* (**B**) in all neuroblastoma cell lines assayed herein. Protein expression for MYCN (**C**) and MYC (**D**) in a subset of cell lines. (**E**) Comparison of unique MYCN peaks called across all MYCN ChIP-Seq samples to known MYCN regulated genes (Valentijn et al. 2012) demonstrates concordance of MYCN ChIP-Seq to MYCN ChIP-ChIP. (**F**) The top 1,330 MYCN peaks (p < 0.05, q < 0.05) are plotted as ChIP-Seq heatmaps for five MYCN amplified cell lines (COG-N-415, Kelly, NB-1643, LAN-5, and NGP) and one MYCN non-amplified cell line (NB-69). All heat map densities ranges from +/-4.0 Kb from the TSS, with average signal plots shown above. (**G**) MYC ChIP-Seq heat maps for the same peaks in two MYCN non-amplified lines (SK-N-AS, and NB-69) and one MYCN amplified cell line (Kelly) showing redundant binding of MYC in non-amplified cell lines.

Next, we evaluated genome-wide binding densities of the histone antibodies and assessed open chromatin by plotting binding of one *MYCN* amplified cell line LA-N-5 (Figure 3A), and one *MYCN* non-amplified cell line NB-69 (Figure 3B). Of note, cell-line specific promoters are located in regions of open chromatin and strongly occupied by narrow regions of H3K4me3 and devoid of H3K27me3 and H3K4me1, as expected. The majority of promoters are also occupied by MYCN in LA-N-5 and MYC in NB-69. Enhancers have bivalent marking of MYCN, H3K4me3, H3K27Ac, open chromatin, and absence of H3K27me3. SEs are broadly marked by MYCN, H3K4me3, H3K27Ac, H3K4me1, and open chromatin.

**Figure 3.**
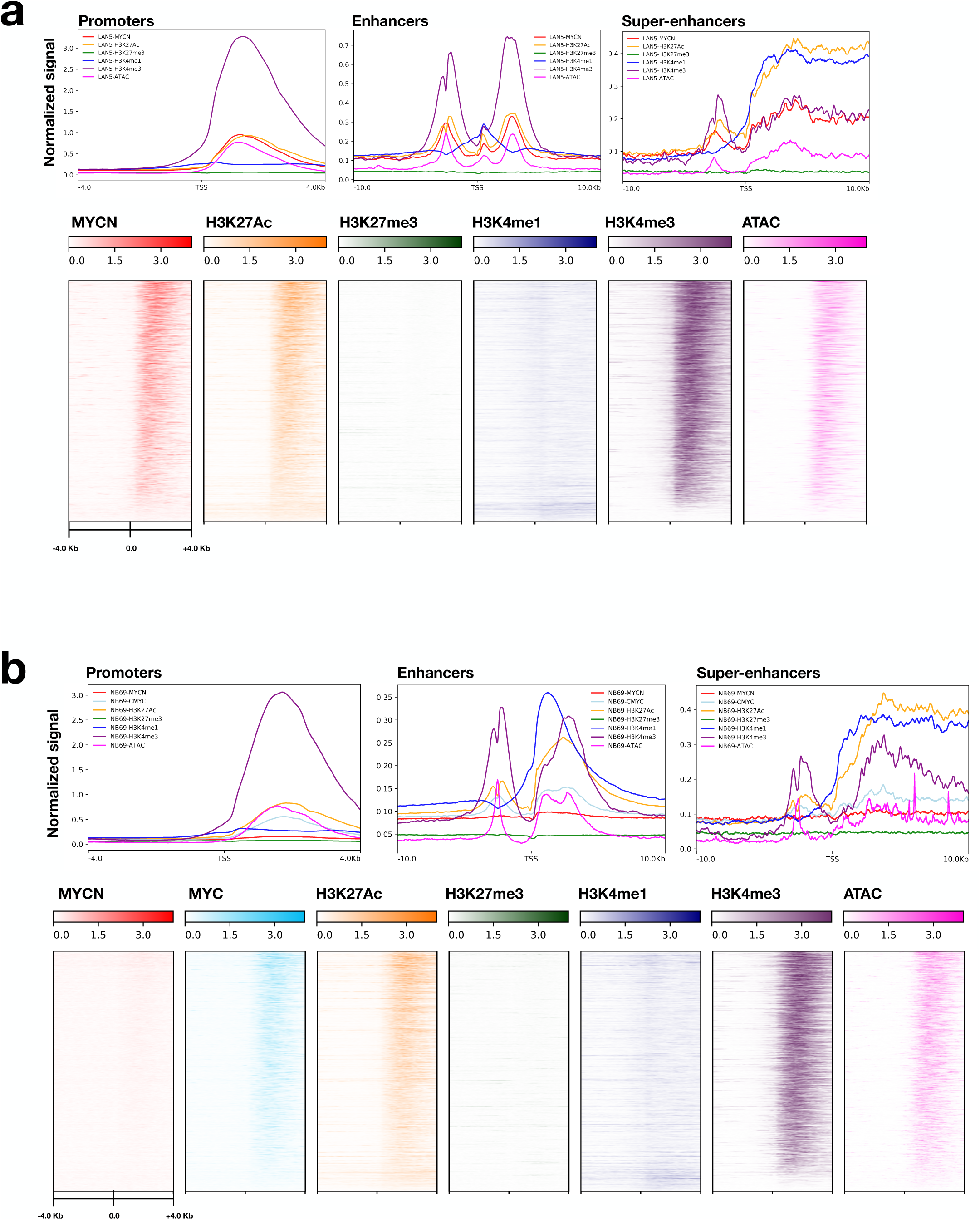
Validation of ChIP-Seq promoter, enhancer, and open chromatin occupancy. (**A**) Binding densities of MYCN, MYC, histone antibodies, and open chromatin for promoter regions of the MYCN-amplified cell line LA-N-5 (**A**) and the MYCN non-amplified cell line NB-69 (**B**) are depicted (+/-4.0 Kb from gene TSS) and distinct profiles are shown for promoters (+/-4.0 Kb from gene TSS), enhancers (+/-10.0 Kb from gene TSS), and SEs (+/-10.0 Kb from gene TSS). For LA-N-5: N_promoters-TSS_ = 4,662, N_enhancers_ = 25,601, N_SE_ = 826, and for NB-69: N_promoters-TSS_ = 4,718, N_enhancers_ = 31,769, N_SE_ = 667.

Finally, we used our H3K27Ac ChIP-Seq data to compare SEs to those reported in two other publications describing the SE landscape in neuroblastoma (Figure 4). Boeva and colleagues identified 5,205 SE-associated genes in Table S3 to identify core regulatory transcriptional circuitry in neuroblastoma using 25 cell lines (Boeva et al. 2017). Four cell lines were common to our study: SK-N-BE(2)C, SK-N-FI, SK-N-AS, and NB-69. Therefore, to validate our H3K27Ac ChIP-Seq, we utilized the same algorithm (LILY, see **Methods**) to call SEs from our H3K27Ac data, and restricted comparison analyses to genes defined as transcription factors (TFs), as defined by core regulatory circuitry (Lambert et al. 2018). We annotated 396 of the SEs reported by Boeva and colleagues as transcription factors and found 59-85% concordance of our TF SE calls (Figure 4A). While a majority of SEs called in each of our cell lines was concordant with Boeva and colleagues, the variance likely stems from the diversity of cell lines in both studies. Thus, we additionally compared our TF SE calls to those from an independent neuroblastoma study (van Groningen et al. 2017) which used the ROSE algorithm (see **Methods**) and reported smaller SE genesets (found in Figure 3) driving the lineage-specific mesenchymal (MES, N = 20 TFs) and adrenergic (ARDN, N = 18 TFs) subtypes. To mimic the analysis performed by van Gronigen and colleagues, we ran ROSE on our H3K27Ac ChIP-Seq data and removed any peaks which overlapped the *MYCN* locus (see **Methods**) to account for false SE calls due to *MYCN* amplification. There were no common neuroblastoma cell lines between van Gronigen and colleagues study and the lines used in our study. We assessed the number of MES or ADRN SE-associated TFs detected in each of our study and found between five and eight ADRN SEs were detected using ROSE (Figure 4B) and between five and 11 ADRN SEs were detected using LILY (Figure 4C). SK-N-SH has a known MES subtype; its subclone, SH-SY-5Y, was profiled as MES by van Gronigen and colleagues. Combining the calls, we were able to significantly (Fisher’s exact test, p < 0.05) validate ADRN subtypes in eight of the ten cell lines we profiled (Figure 4D). Interestingly, SK-N-AS contains SEs from both subtypes and thus may reflect a heterogeneous cell line. Specific SEs are reported per algorithm per cell line in Online Table 5. As further validation, we re-analyzed publicly-available SK-N-SH H3K27Ac (Biosample SAMN05733860, Run SRR5338927) and SK-N-SH Input (Biosample SAMN05733844, Run SRR5471111) ChIP-Seq data (GEO accession GSM2534162) using the same peak-calling and SE pipelines used on our data (see **Methods**). We observed enhancer binding (H3K27Ac) and open chromatin (ATAC) at the same loci we observe strong MYC occupancy (Supplemental Figure 1A). Further, we assessed concordance of SEs called in SK-N-SH with those previously reported and found 76% of TF SEs called in SK-N-SH in common with those from Boeva, et. al, similar to our findings.

**Figure 4.**
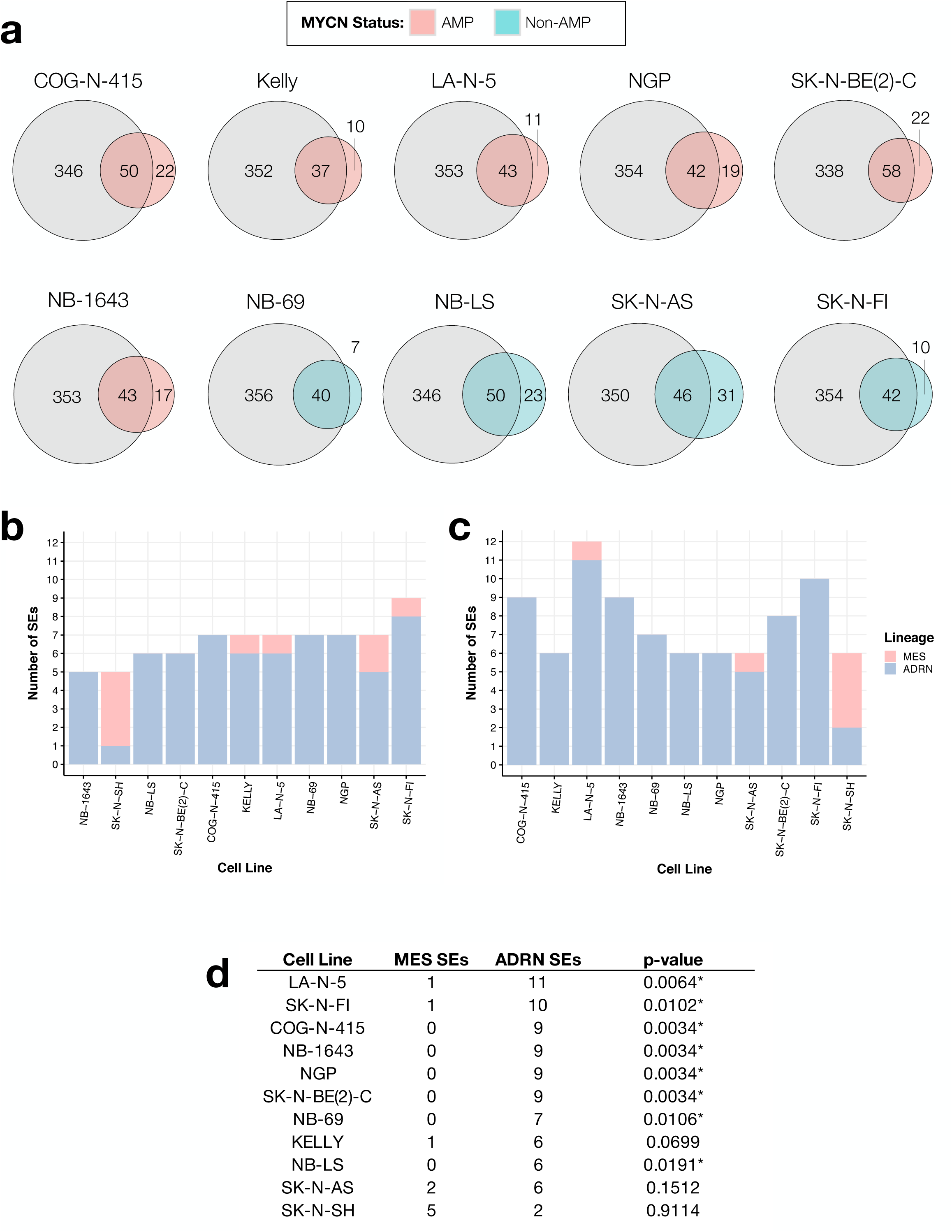
Comparison of super-enhancers called to literature reports. SEs were called using the H3K27Ac ChIP-Seq data collected in ten neuroblastoma cell lines plus SK-N-SH H3K27Ac ChIP-Seq data from ENCODE. Results were compared to two other literature sources which examined SE signatures in neuroblastoma. (**A**) LILY TF-specific SEs were 59-85% concordant with (Boeva et al. 2017). (**B**) Total number of lineage-specific SEs called from ROSE and (**C**) LILY per cell line (ADRN = adrenergic, MES = mesenchymal subtype). (**D**) Total number of lineage-specific SEs called when combining ROSE and LILY results. One-tailed fisher’s exact test p-values are listed (*denotes significance at an α level of 0.05).

Together, we have validated both MYCN and MYC ChIP-Seq antibodies for use in ChIP-Seq, as well as genome-wide occupancy profiles for histone markers and open chromatin across a cohort of neuroblastoma cell lines. We ran two algorithms (LILY and ROSE) and compared our data to two independent datasets to validate reproducibility of lineage-specific SEs in neuroblastoma cell lines. Finally, we demonstrate integration of publicly-available H3K27Ac data from SK-N-SH with our MYC ChIP-Seq and ATAC-Seq data, and show reproducibility of SE calls between the publicly-available data and two independent reports. These data should be a valuable resource to the childhood cancer and MYC research communities.

## Usage Notes

Here, we provide a comprehensive, validated ChIP-Seq (MYCN, MYC, H3K27Ac, H3K27me3, H3K4me3, and H3K4me1) and ATAC-Seq neuroblastoma cell line dataset which can be coupled with our previous RNA-Seq profiling dataset (Harenza et al. 2017) to interrogate novel transcriptional regulation in this disease. For example, the H3K27me3 ChIP-Seq can be used to identify genes being repressed via the PRC2 complex, while H3K27Ac and H3K4me1 ChIP-Seq can be used to interrogate promoter-enhancer mechanisms. CSI-ANN can be used to integrate histone ChIP-Seq data to predict regulatory DNA segments (Firpi, Ucar, and Tan 2010), and IM-PET can use the results from CSI-ANN to predict enhancer-promoter interactions without the need for Hi-C data (He et al. 2013). Additionally, chromatin states can be inferred (Ernst and Kellis 2012; Sohn et al. 2015), and these data can be later integrated with whole exome or genome sequencing data or genome-wide association studies to identify molecular alterations driving transcriptional regulatory marked by histone marks or open chromatin.

All data are openly-available from GEO as described in the Data Records section.

## Supporting information

MetadataTable1

OnlineTable1

OnlineTable2

OnlineTable3

OnlineTable4

OnlineTable5

## Code Availability

Code for SE calling, filtering, and heatmap generation is available on Github (https://github.com/marislab/epigenomics-data-descriptor).

## Acknowledgements

Funding for this research was provided by NIH grants R01 CA180692 (JMM), R35 CA220500 (JMM), and an Alex’s Lemonade Stand Young Investigator Award (JLR). We thank the neuroblastoma patients and families for donating tumor tissue from which cell lines used in this study were derived.

## Author contributions

Conceptualization: JLR, KU

Methodology: JLR, KU, KP, RTS, RNA, PF, GPW

Validation: JLR, KU, KP, KLC, AM

Formal Analysis: JLR, KU, KP, AM

Investigation: JLR, KU, KP

Resources: JMM, SJD

Data Curation: JLR, KP, GIS

Writing -Original Draft: JLR, KU, KP, AM

Writing -Review & Editing: JLR, KLC, KU, GPW, JMM

Visualization: JLR, KU, KP

Supervision: JLR

Funding Acquisition: JLR, JMM

## Table Legends

**Online Table 1. Neuroblastoma cell lines profiled in this study.**

Identification of which cell lines were used to perform ChIP-Seq and ATAC-Seq, along with their MYCN amplification status.

**Online Table 2. Materials and reagents used in this study.**

Information regarding the resources used for obtaining the ChIP-Seq and ATAC-Seq data. This includes manufacturer, item number or identification code, and the volume or concentration used.

**Online Table 3. ATAC-Seq primer sequences.**

Primer sequences used in ATAC-Seq to amplify transposed DNA.

**Online Table 4. ChIP-Seq Quality Control Metrics.**

Total number of reads, FRiP score, RSC, and NSC per sample.

**Online Table 5. Super-enhancer-associated transcription factor comparisons.**

## Manuscript metadata tables

**Metadata Table 1. Neuroblastoma cell line metadata.**

Listed are the cell lines used in this study, MYCN amplification status, and culturing media information.

---

**Supplemental Figure 1.**
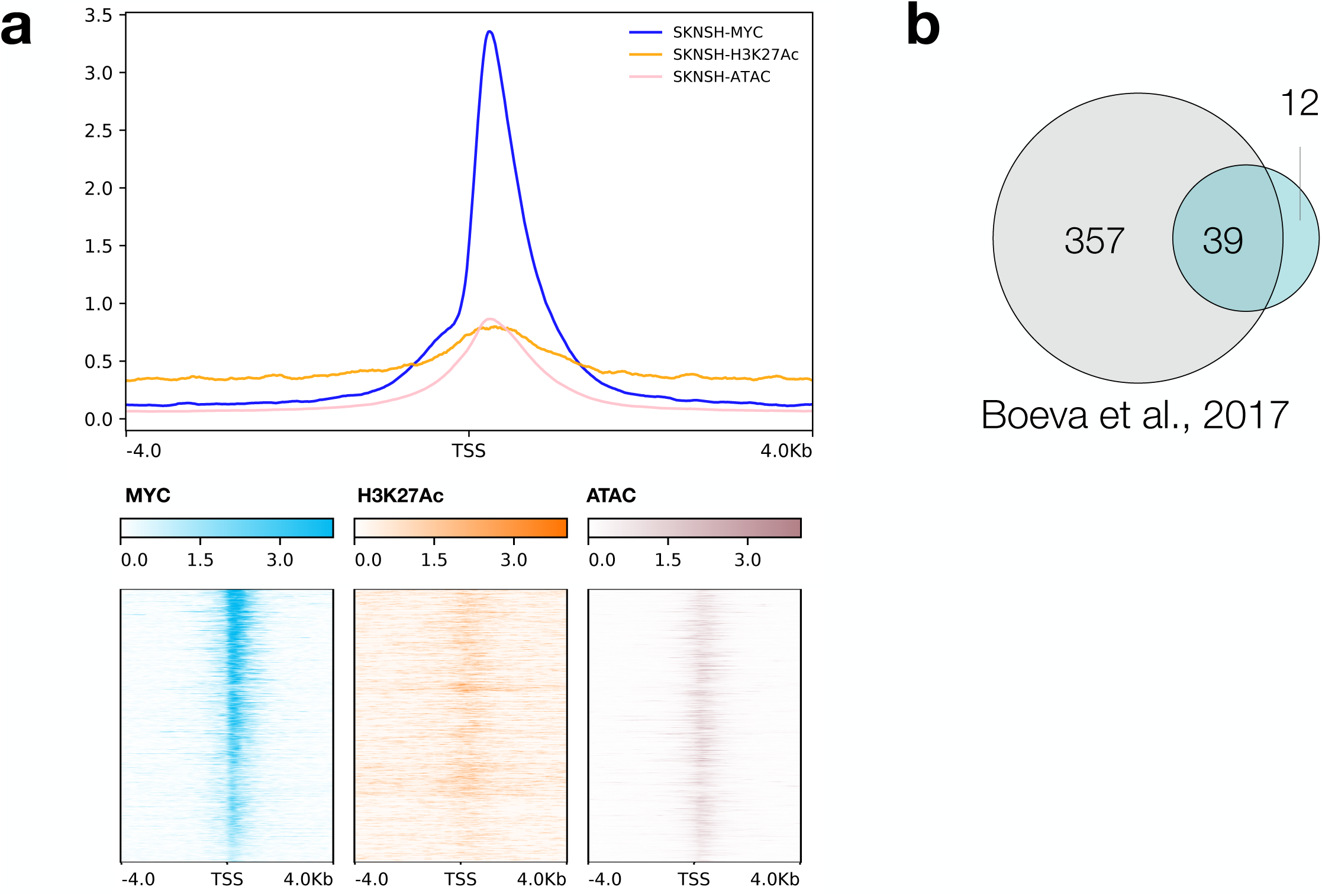
Integration of ENCODE SK-N-SH H3K27Ac ChIP-Seq with our dataset. Depicted are heatmaps and the binding profiles **(a)** for SK-N-SH MYC, H3K27Ac (ENCODE), and ATAC-Seq for the top 5,000 MYC peaks. **(b)** LILY super-enhancer calls were 76% concordant with Boeva, et. al, 2017 (N = 39/51).

